# Dimensions of passerine biodiversity along an elevational gradient: a nexus for historical biogeography and contemporary ecology

**DOI:** 10.1101/842138

**Authors:** Kevin R. Burgio, Steven J. Presley, Laura M. Cisneros, Katie E. Davis, Lindsay M. Dreiss, Brian T. Klingbeil, Michael R. Willig

## Abstract

**Aim:** The incorporation of functional and phylogenetic information is necessary to comprehensively characterize spatial patterns of biodiversity and to evaluate the relative importance of ecological and evolutionary mechanisms in molding such patterns. We evaluated the relative importance of mechanisms that shape passerine biodiversity along an extensive elevational gradient.

**Location:** Manu Biosphere Reserve in the Peruvian Andes

**Taxon:** Songbirds (order Passeriformes)

**Methods:** We quantified elevational gradients of species richness, phylogenetic biodiversity, and functional biodiversity for all passerines as well as separately for suboscines and oscines; determined if phylogenetic or functional biodiversity was consistent with random selection or if there was evidence of particular mechanisms dominating community assembly; and compared patterns for each dimension of biodiversity for the two suborders.

**Results:** For all passerines and for suboscines, species richness decreased in a saturating fashion, phylogenetic biodiversity declined linearly, and functional biodiversity was stochastic along the elevation gradient. For oscines, species richness and phylogenetic biodiversity decreased linearly, and functional biodiversity decreased in a saturating fashion.

**Main conclusions:** Elevational gradients of biodiversity at Manu result from a combination of adaptations associated with radiations that occurred elsewhere (suboscines in Amazonian lowlands, oscines in colder climes of North America) and an in situ radiation in the Andes (tanagers). Our results suggest a combination of temperature-related physiological constraints and a reduction in functional redundancy associated with decreasing resource abundance at higher elevations molded the passerine assemblages along this elevational gradient. Explicit consideration of historical biogeography and conservatism of ancestral niches is necessary to comprehensively understand the mechanisms that mold gradients of biodiversity.

## INTRODUCTION

During the first part of the 21^st^ century, the quantification of spatial patterns of biodiversity improved the identification of ecological and evolutionary processes that structure communities at multiple spatial scales. Such understanding, which is vital to accurately forecast the effects of environmental change on ecological communities, requires exploring the diversity of species functions and evolutionary history (i.e. genetic potential). Despite the increased use of functional and phylogenetic information to understand spatial patterns of biodiversity and mechanisms of community assembly, conservation policies often focus on biotas that represent a single evolutionary radiation or that represent a single functional guild (e.g. Devictor et al. 2010, Cisneros et al. 2014, Dehling et al. 2014, Dreiss et al. 2015). More recently, incorporation of multiple dimensions of biodiversity has increasingly characterized conservation planning (e.g. Mazel et al. 2014, Kosman et al. 2019, Burgio et al. 2019).

The phylogenetic dimension of biodiversity (PD) reflects variation in evolutionary history among species and is based on time since divergence from a common ancestor (Vellend et al. 2010). Because of the broad and rapid changes in climate and land use occurring now and predicted to continue in the near future, the capacity for future adaptation may be an important aspect of biodiversity to consider for effective medium- and long-term conservation (Cardinale et al. 2012). The functional dimension of biodiversity (FD) reflects variation in the ecological attributes of species. Traits that quantify FD of an assemblage can provide insight about functional uniqueness, redundancy, or complementarity (Walker 1992, Vandewalle et al. 2010), and may better represent the degree of maintenance, resistance, or resilience of ecosystem services in the face of species loss. Importantly, each of these dimensions can offer information complementary to that provided by taxonomic considerations. Consequently, incorporating taxonomic, phylogenetic, and functional perspectives into evaluations of spatial patterns of biodiversity is necessary to understand mechanisms that give rise to spatial patterns (Presley et al. 2018, Montaño-Centellas et al. 2019) and to forecast the efficacy of conservation options (Burgio et al. 2019).

Major ecological theories for biodiversity patterns along elevational gradients are similar to those for latitudinal gradients (Willig and Presley 2016): species richness is generally thought to decline with increasing elevation for a number of reasons, including decreasing resource abundance and diversity (Jankowski et al. 2013), and environmental factors that result in species filtering at higher elevations (e.g. lower temperatures and increased seasonality; Coyle et al. 2014). Competition is another mechanism that is used to explain elevational diversity gradients, though views on its effect are conflicting: an increase in interspecies competition may lead to local extinctions and decreased diversity (Graham et al. 2014) or niche partitioning and increased diversity (Cavendar-Bares et al. 2009, Burns and Strauss 2012). These conceptual frameworks have been expanded to develop and employ analytical approaches to estimate biodiversity based on evolutionary histories or ecological functions of species in an assemblage (e.g. Webb et al. 2002, Pavoine and Bonsall 2011). In particular, frameworks for evaluating the extent to which variation in phylogenetic or functional biodiversity is a consequence of variation in species richness (S) provide further insight into the relative importance of various processes (e.g. abiotic or biotic filtering, niche partitioning, interspecific competition) in structuring communities along gradients (e.g. Lopez et al. 2016).

In general, PD or FD is expected to increase in a saturating fashion with increasing species richness because the probability of adding species with novel ecological attributes or that represent a different evolutionary lineage to an assemblage decreases as assemblages become more species rich (Kluge and Kessler 2011). Relationships between FD and species richness can determine the degree to which local community assembly is dominated by niche differentiation and competition, or by adaptation to the local environment (Kluge and Kessler 2011). Alternatively, relationships between PD and species richness can determine the degree to which competition within or between clades dominates local community assembly (Mayfield and Levine 2010). Because trait conservatism is commonly strong for important functional traits (e.g. Ackerly 2003, Wiens et al. 2010), spatial patterns of phylogenetic and functional biodiversity are typically highly correlated or qualitatively similar, resulting in similar conclusions for each of these two dimensions (e.g. Cadotte et al. 2009, Devictor et al. 2010, Flynn et al. 2011, Meynard et al. 2011, Cisneros et al. 2014, Dehling et al. 2014, Dreiss et al. 2015). This may occur because traits and functions of organisms arise through modification of those inherited from ancestors (Shanahan 2011), which gives rise to the a priori expectation that more closely related species will be functionally more similar (Safi et al. 2011) and would result in a positive relationship between PD and FD. Nonetheless, comparison of how the two dimensions differ along an elevational gradient can be used to infer community assembly processes.

If PD or FD of a local community is indistinguishable from that of randomly selected sets of species from the regional pool, community assembly may be random with respect to functional or evolutionary characteristics of species. PD greater than expected by chance indicates that within-clade competition dominates assembly processes, whereas PD less than expected by chance is consistent with inter-clade competition dominating assembly (Mayfield and Levine 2010). FD greater than expected by chance suggests that some combination of niche partitioning, limiting similarity, and character displacement dominates community assembly, whereas FD less than expected by chance suggests that environmental filtering (abiotic or biotic) structures communities (Kluge and Kessler 2011). Finally, if processes that have opposing effects are relatively equal in strength, communities will be indistinguishable from those structured by random selection. In addition to evaluating deviations from expectations along a spatial gradient, one can identify systematic changes in the relative importance of ecological or historical factors in affecting the composition of local communities.

For birds, PD and FD generally decrease with elevation after controlling for variation in species richness, although there is a great deal of variation among elevational gradients (Montano-Centallas et al. 2019). However, at the scale of all bird orders, important patterns may be obscured, especially patterns informed by different biogeographic histories of particular clades. For instance, evolutionary history and biogeographic radiations of the two suborders of Passeriformes likely played a role in shaping the PD and FD differences among various habitats in Brazil (Almeida et al 2018).

Passerines are among the most species-rich, abundant, and functionally diverse groups of vertebrates (Barker et al. 2004). The Passeriformes include two major suborders: the Passeri or oscines, which are the true songbirds, and the Tyranni or suboscines. The two groups diverged between 77 mya (Barker et al. 2004) and 60 mya (Selvatti et al. 2015), largely coinciding with the split of Australia and eastern Antarctica. Suboscines arrived and diversified in South America ~64 mya, prior to the principal orogeny of the Andes while oscines radiated multiple times prior to their arrival in South America ~47 mya (Barker et al. 2004). Despite their distinctly different evolutionary histories, foraging strategies, and physiologies, each suborder exhibits similar functional breadth comprising many of the same foraging guilds (e.g. aerial insectivores, ant-following insectivores, and frugivores), making them an excellent model for understanding the effects of environmental variation on community structure and biodiversity. The arrival, radiation, and diversification of passerines along the extensive environmental gradient represented by the Andes make this system particularly suitable for exploring large-scale patterns in multiple dimensions of biodiversity.

In this study, we evaluated species richness (S), phylogenetic biodiversity, and functional biodiversity along an extensive elevational gradient for a passerine fauna with a complex history of evolutionary radiations. We used data from the Manu Biosphere Reserve in the Peruvian Andes to comprehensively explore differences between these evolutionary radiations to better understand the relative importance of mechanisms that structure these passerine assemblages. We hypothesized that suboscines, which diversified in the tropical forests of South America would have “incumbency advantage” in the hotter lower elevations, whereas oscines, which radiated into Peru through North America, would have the advantage in cooler, upper elevations. Because of “incumbency advantage” we expected suboscines would have higher levels of S and FD in the lower elevations, but due to the more complex biogeographic history of the oscines, we expected them to have higher FD and PD at the upper end of the gradient. To test this, we (1) described the elevational gradients of S, PD, and FD for all passerines, as well as for suboscine and oscine components separately; (2) determined if PD or FD for each taxon was consistent with random selection or if they show evidence of particular mechanisms dominating community assembly; and (3) compared species richness, PD, and FD of suboscines and oscines, clades with different evolutionary histories and functional ecologies, to determine changes in the relative contributions of each clade to biodiversity of local assemblages.

## MATERIALS AND METHODS

### Study area and organisms

Manu, a UNESCO Biosphere Reserve (11°51′23″S, 71°43′17″W) and an IUCN World Heritage site, is one of the most diverse protected tropical areas in the world (MacQuarrie 1992, Patterson et al. 2006). Located along the eastern slopes of the Andes of southeastern Peru, the elevational gradient of Manu (340 to 3625 m above sea level) supports diverse faunal and floral assemblages, including 1005 bird species (Walker et al. 2006). Structurally distinct vegetation types replace one another along the gradient (Terborgh 1971, Patterson et al. 1998). Lowland rainforest occurs below 500 m and is characterized by 50 to 60 m canopies. Montane rainforest occurs from 500 to 1400 m, and is less vertically complex (~ 35 m tall), with a less well defined subcanopy. A persistent cloud layer results in cloud forest from 1400 to 2800 m. Elfin forest extends from 2800 to 3200 m, and supports a low canopy (~ 15 m) and dense vegetation. Above 3200 m, patches of elfin forest are intermixed with tall grasslands.

We compiled comprehensive data from museum specimens, published literature, and recent surveys on the elevational distributions of 584 species of passerines from Manu, excluding vagrants (10 species) and migrants (40 species) (Walker et al. 2006). Species incidence data (i.e. presence or absence) were organized into thirteen elevational intervals that each span 250 m, with the upper boundaries defining stratum names (e.g. the elevation range from 250 m to 500 m is the 500 m stratum). Several expeditions were conducted from 1997–2001 to address historically uneven sampling effort along the gradient, which focused on mid- and high-elevations and comprised six person-years of effort (Patterson et al. 2006). Those expeditions resulted in the addition of 44 species of passerine to the list of known species from Manu, many of which are mid- or high-elevation specialists. The completeness of the passerine data due to the extensive, intensive, and long-term nature of surveys at Manu is corroborated by the existence of no remaining gaps in the elevational distributions of the 534 resident species, and the extraordinary level of coherence for this metacommunity (Presley et al. 2012). To explore contemporary ecological and historical biogeographical mechanisms driving elevational gradients of biodiversity, we quantified the dimensions of biodiversity at each stratum, and conducted analyses separately for all passerines (534 species), for oscines (206 species), and for suboscines (328 species). We followed the nomenclature and taxonomic recommendations of the South American Classification Committee of the American Ornithologists’ Union (Remsen et al. 2012).

### Quantification of dimensions of biodiversity

We estimated PD and FD by Rao’s quadratic entropy (Rao’s Q, Botta-Dukát 2005). Rao’s Q measures average interspecific difference between species, thereby reflecting multivariate dispersion. Average phylogenetic or functional distances were obtained from a pairwise dissimilarity matrix for the phylogenetic component and for each of five functional approaches. To promote meaningful comparisons among dimensions of biodiversity, we transformed Rao’s Q into its effective number of species or Hill number (hereafter numbers equivalent). The numbers equivalent is the number of maximally dissimilar species that is required to produce the empirical value of a metric (Jost 2006). This transformation facilitates intuitive interpretation of differences among assemblages and dimensions because indices are expressed in the same units (Jost 2006). Species richness is its own numbers equivalent. We quantified Rao’s Q as a numbers equivalent using R functions (de Bello et al. 2010) for PD and FD.

We estimated phylogenetic differences using branch lengths derived from a comprehensive phylogeny of all birds, which included all of the passerines from Manu (Jetz et al. 2012). This phylogeny was inferred using a two-step protocol in which time-calibrated phylogenetic trees were estimated for well-supported bird clades and subsequently joined onto a backbone tree representing deep phylogenetic relationships. This method allows for species-level inference that also reflects uncertainty. We used the phylogeny subset tool from birdtree.org to download a distribution of 1000 trees containing only the passerine species of Manu. Trees were based on the “Hackett All Species” pseudo posterior distribution. For each of the 1000 phylogenetic trees, we calculated a pairwise dissimilarity matrix via the “cophenetic” function of the R package “ape” (Paradis et al. 2004), and we estimated PD as the mean of Rao’s Q values calculated from each of the 1000 dissimilarity matrices.

We estimated functional differences among species using two types of data: categorical (binary) and mensural attributes (Table 1; Supplementary material Table S1). Subsets of these functional attributes defined particular functional components. Categorical attributes represented diet, foraging strategy, or foraging location, traits commonly used in gradient studies (e.g. Cisneros et al. 2014, Dreiss et al. 2015, Montano-Centellas et al. 2019). Mensural attributes were represented by body size, a surrogate for environmental tolerance and caloric requirements. For each categorical attribute, a species received a “1” if it exhibited the characteristic or a “0” if it did not. For each mensural attribute, we obtained average values based on the measurement of adult individuals from the literature (Supplementary material Table S1). We derived information for all attributes from the literature or museum records, and restricted records to those from South America when possible (Supplementary material Table S1). We estimated missing mensural values via linear regressions of mass and length using attribute values of other species from the same genus or family. For missing categorical data, we substituted attributes from congeners. We substituted only 1.2% (131 of 10,680) of species traits.

**Table 1:** Functional attributes that reflect niche axes (functional components) were τsed to estimate functional biodiversity of passerine assemblages at Mam Mensiral attributes were measured as described in sources (see table SI, Supporting Information).

| Type of Data | Functional Component | Attribute | Trait Values |
| --- | --- | --- | --- |
| Categorical | Diet | Vertebrates | 1, 0 |
|  |  | Invertebrates | 1, 0 |
|  |  | Nectar | 1, 0 |
|  |  | Seed | 1, 0 |
|  |  | Fruit | 1, 0 |
|  | Foraging Strategy | Pursuit | 1, 0 |
|  |  | Glean | 1, 0 |
|  |  | Pounce | 1, 0 |
|  |  | Dig | 1, 0 |
|  |  | Scavenge | 1, 0 |
|  |  | Follow Ants | 1, 0 |
|  |  | Flower Pierce | 1, 0 |
|  |  | Probe | 1, 0 |
|  | Foraging Location | Water | 1, 0 |
|  |  | Ground | 1, 0 |
|  |  | Vegetation | 1, 0 |
|  |  | Dead Leaves | 1, 0 |
|  |  | Air | 1, 0 |
| Mensural | Body Size | Mass | Mean (g) |
|  |  | Length | Mean (cm) |

To best portray the variety of functions performed by a species, we considered all attributes of a particular functional component (e.g. diet or body size) together in defining that functional component. Particular functional components may differ in the way they respond to environmental gradients. In such circumstances, integration of all ecological attributes into a single multivariate measure may obscure important patterns (Spasojevic and Suding 2012). Thus, we estimated functional differences among species for each functional component separately (Table 1), as well as for all functional components combined. For analyses based on all functional components, we weighted each component equally to account for variation in the number of attributes associated with components. In all cases, we based values of Rao’s Q on functional dissimilarity matrices calculated using the Gower metric from the R package “cluster” (Maechler et al. 2012).

### Quantitative analyses

To discriminate among different elevational relationships (e.g. random, linear, and nonlinear relationships such as saturating, modal, or u-shaped), we evaluated the significance of parameters associated with a second-order polynomial (Dutka and Ewens 1971) for each combination of group (i.e. passerines, oscines, and suboscines) and dimension of biodiversity. We used the family of second-order polynomials to capture linear and non-linear responses of different dimensions of biodiversity to elevational variation, as we had no *a priori* expectation or theoretical argument to support the exploration of higher-order polynomials. We used orthogonal regression because it decomposes the general relationship from ordinary polynomial regression to provide independent and additive estimates of the importance of magnitude (b^*^_0_), a constant rate of change (b^*^_1_), and a varying rate of change (b^*^_2_) to elevational gradients. Also, we conducted one-tailed Spearman rank correlations to determine if elevational gradients of PD, FD, and each component of FD were positively correlated with variation in species richness.

To evaluate the extent to which the elevational relationship of PD, FD, or components of FD arose as a consequence of the random selection of species from the regional pool, we conducted a suite of randomizations using the trial-swap method (Miklós and Podani 2004). This method randomizes the phylogenetic or functional identities of species at each elevational stratum while constraining species richness to equal its empirical value and constraining the frequency of occurrence for species during the simulations to equal empirical frequencies along the gradient (The R package “picante,” Kembel et al. 2010). The composition of each species pool corresponded to species identities in each taxonomic group at Manu (i.e. passerines, oscines, or suboscines), regardless of elevation. We executed 1000 iterations for each analysis of FD, including analyses conducted separately for each component of FD, and 1000 iterations for each of the 1000 phylogenetic trees for considerations of PD. For each iteration, we calculated *b_1_^*^* and *b_2_^*^* using orthogonal polynomial regression, thereby creating a distribution of 1000 values for each regression coefficient for FD and 1,000,000 values for each regression coefficient for PD. We considered an empirical coefficient to be different from those generated by chance if it was greater than or less than 97.5% of the values generated via simulation (twotailed test).

Separately for each dimension and each component of FD, we compared the elevational gradients for oscines and suboscines via linear (b_1_) and non-linear (b_2_) parameters from ordinary polynomial regression as well as via the coefficients that represent independent contributions of linear (b^*^_1_) and non-linear (b^*^_2_) components from orthogonal polynomial regression. If confidence intervals (i.e., ± 2 SE) for analogous parameters did not overlap between suboscines and oscines, they were considered to be significantly different. In addition, we used paired t-tests for each dimension separately to determine if the magnitude of elevation-specific diversity of one suborder was consistently different from that of the other.

## RESULTS

Elevational gradients of biodiversity were distinct for each dimension (Fig. 1–2). For all passerines and suboscines, species richness declined with elevation in a saturating fashion, with a large proportion of the variation in species richness (R^2^ = 0.94 and R^2^ = 0.95, respectively) related to variation in elevation (Supplementary material Tables S2 and S3). For oscines, species richness declined linearly (R^2^ = 0.91). For each taxonomic group, PD declined linearly with elevation, with no evidence of saturation at high elevations (Fig. 2), with the model accounting for 87.8%, 82.3%, and 60.0% of the variation in PD for passerines, suboscines, and oscines, respectively. In contrast, elevational variation in FD was stochastic for all passerines and suboscines, with non-significant linear and quadratic components, as variation in elevation accounted for only 5% and 16% of variation in FD, respectively (Fig. 2). In contrast, elevational variation in FD of oscines had significant linear and quadratic components, with elevational variation accounting for 78% of the variation in FD. Importantly, for each functional component, oscines and suboscines evinced different responses along the elevational gradient (Fig. 3, Supplementary material Table S3).

**Figure 1.**
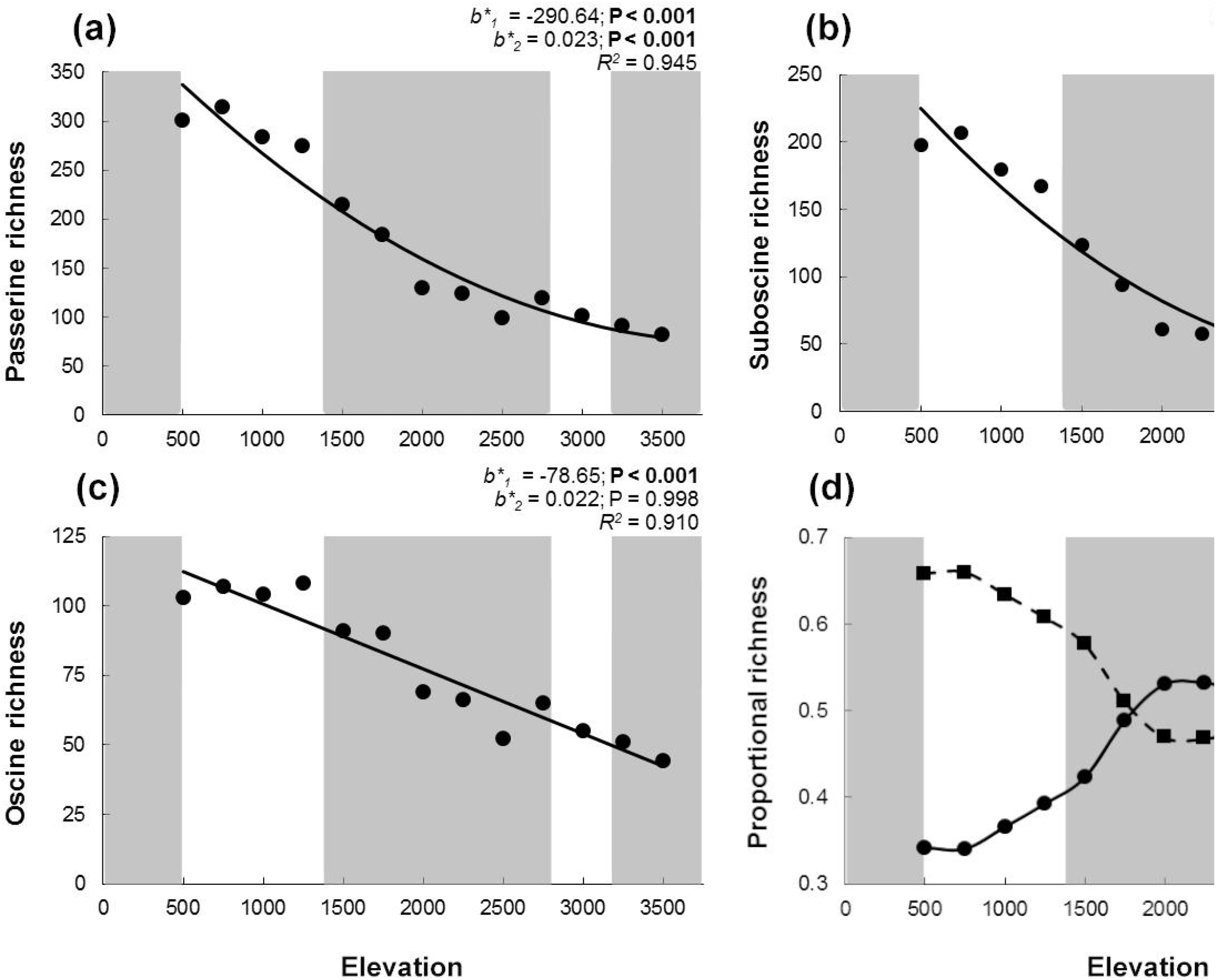
Elevational variation in species richness for all passerines (a), suboscines (b), and oscines (c) at Manu. Empirical values are represented by black dots. A solid line represents an empirical polynomial relationship, and adjusted R^2^ is the fit of the model. Significant (P ≤ 0.05) orthogonal regression coefficients (b*1, and b*2) are bold. Elevational variation of the proportional richness of suboscines (squares) versus oscines (circles) documents a change in suborder dominance at mid-elevations (d). Alternating shaded regions of the graphic correspond to elevationally defined forest types: lowland rainforest, montane rainforest, cloud forest, elfin forest, and mixed elfin forest-grassland.

**Figure 2.**
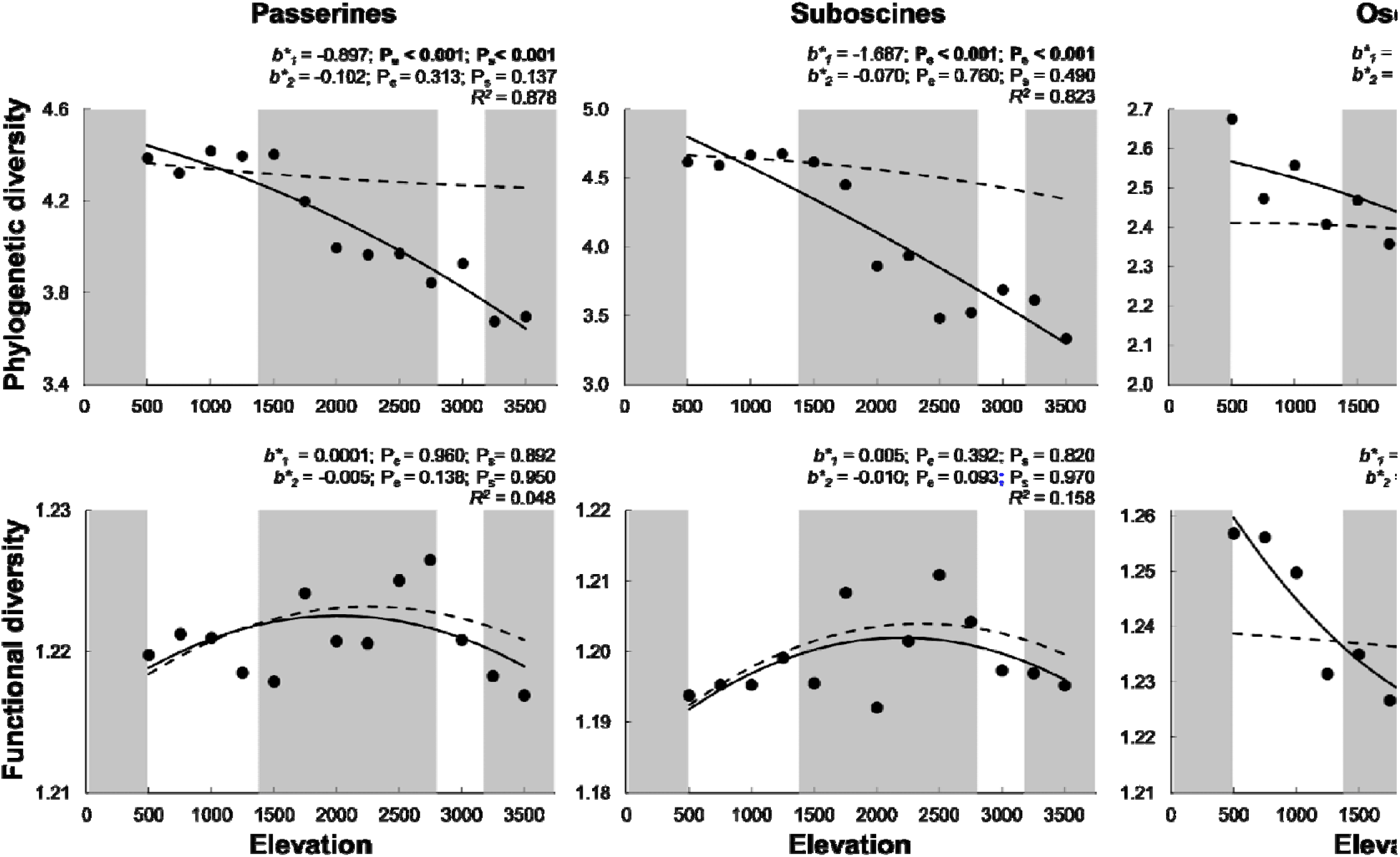
Elevational gradients of phylogenetic diversity and functional diversity for all passerines, suboscines, and oscines. Empirical values of Rao’s Q (transformed as Hill numbers) are represented by black dots. A solid line represents the empirical polynomial relationship and adjusted *R*^2^ is the fit of the model. Significant orthogonal regression coefficients (*b^*^_1_*, and *b^*^_2_*) for empirical gradients are bold (*P_e_* ≤ 0.05). Dashed lines represent mean expected polynomial relationships based on random sampling. Significant differences (*P_s_* ≤ 0.05) between empirical orthogonal regression coefficients and randomly generated orthogonal regression coefficients are indicated by bold. Alternating shaded regions of the graphic correspond to elevationally defined forest types: lowland rainforest, montane rainforest, cloud forest, elfin forest, and mixed elfin forest-grassland.

**Figure 3.**
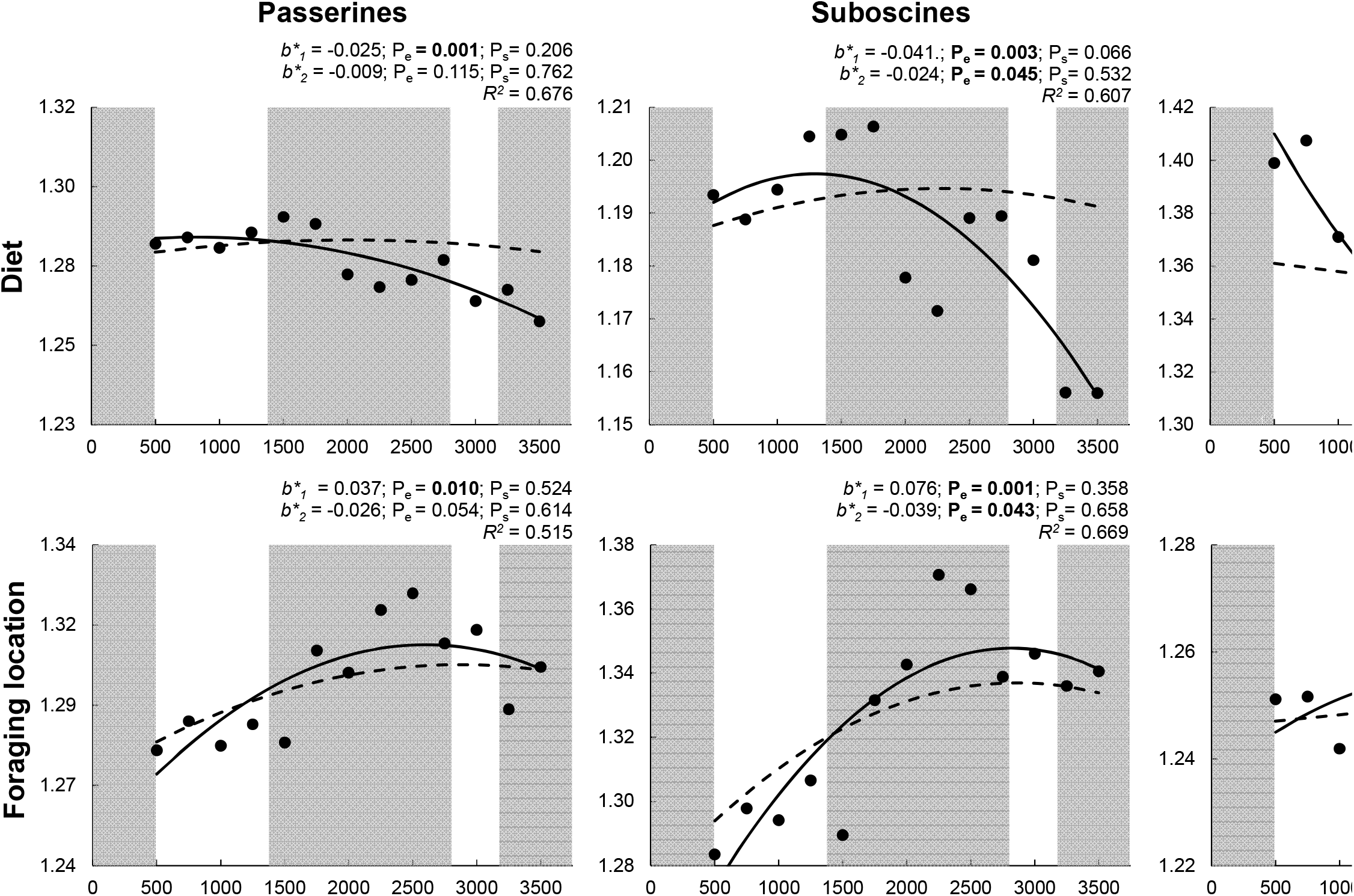
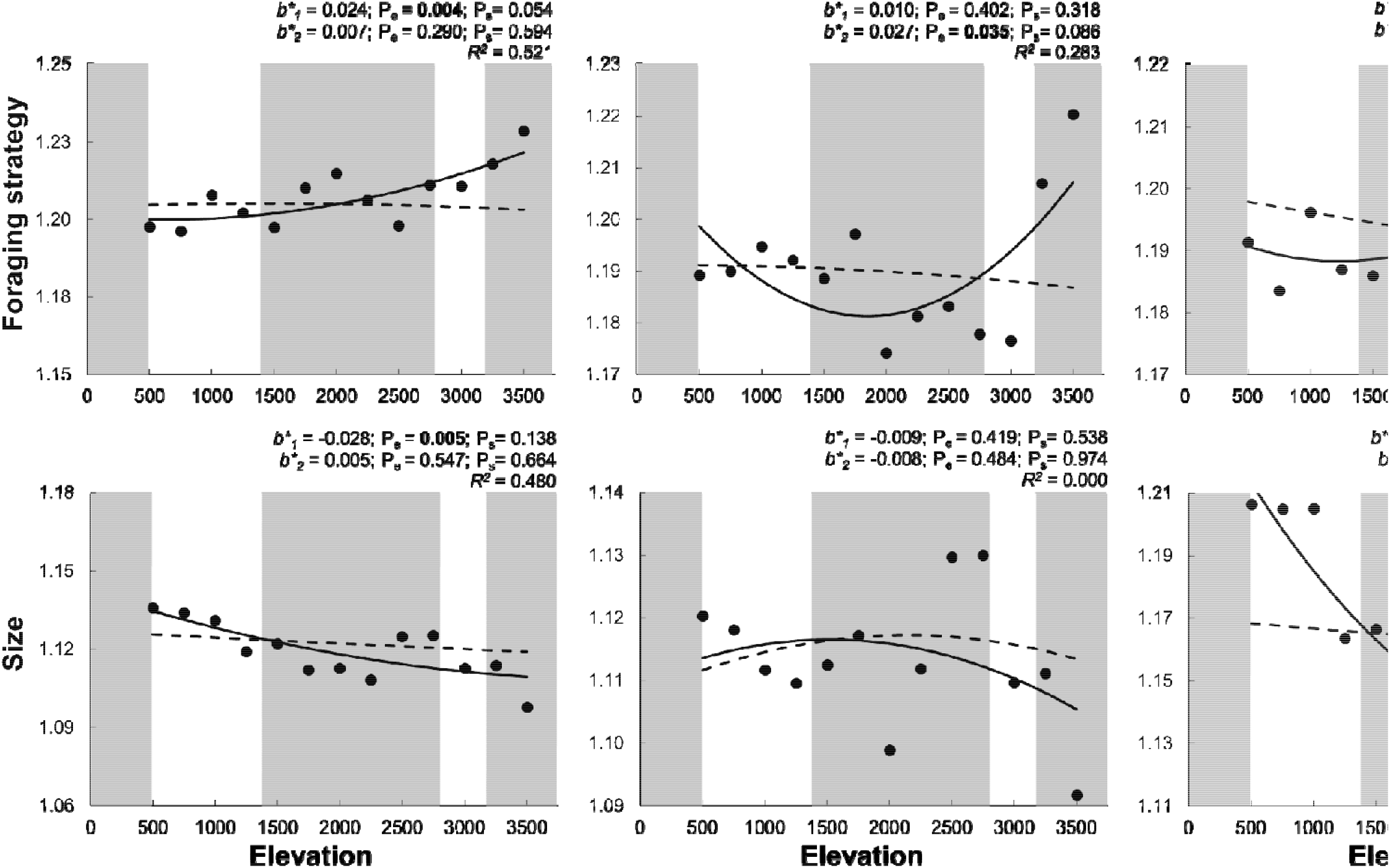
Elevational gradients for each of four components of functional diversity for all passerines, suboscines, and oscines. Empirical values of Rao’s Q (transformed as Hill numbers) are represented by black dots. A solid line represents the empirical polynomial relationship, and adjusted *R* is the fit of the model. Significant orthogonal regression coefficients (*b^*^_1_*, and *b^*^_2_*) for empirical gradients are bold (*P_e_* < 0.05). Dashed lines represent mean expected polynomial relationships based on random sampling. Significant differences (*P_s_* < 0.05) between empirical orthogonal regression coefficients and randomly generated orthogonal regression coefficients are bold. Alternating shaded regions of the graphic correspond to elevationally defined forest types: lowland rainforest, montane rainforest, cloud forest, elfin forest, and mixed elfin forestgrassland.

Species richness was correlated positively with PD for each taxonomic group but was only positively correlated with FD for oscines (Table 2). Species richness was correlated positively with diet diversity for each group, and with size diversity for all passerines and oscines. Species richness was not correlated positively with foraging location or foraging strategy diversity for any group (Table 2).

**Table 2.** Spearman rank correlations between species richness and phylogcnclic diversity, functional diversity, or components of functional diversity for all passerines, for suboscines, or for oscines at Mann Biosphere Reserve. Significant (P ≤ 0.05) are bold. Analyses were conducted as one-tailed tests because expectations are that dimensions of biodiversity should be positively associated with species richness.

|  | All passerines |  | Suboscines |  | Oscines |  |
| --- | --- | --- | --- | --- | --- | --- |
| | $\rho$ | P-value | $\rho$ | P-value | $\rho$ | P-value |
| Phylogenetic diversity | **0.863** | **< 0.001** | **0.874** | **< 0.001** | **0.560** | **0.023** |
| Functional diversity | 0.126 | 0.341 | -0.269 | 0.818 | **0.714** | **0.004** |
| Diet diversity | **0.786** | **0.001** | **0.632** | **0.012** | **0.720** | **0.004** |
| Foraging location diversity | -0.687 | 0.994 | -0.725 | 0.997 | 0.060 | 0.425 |
| Foraging strategy diversity | -0.720 | 0.997 | 0.011 | 0.489 | -0.456 | 0.943 |
| Size diversity | **0.632** | **0.012** | 0.313 | 0.149 | **0.758** | **0.002** |

PD declined with elevation more quickly than expected based on random selection for each taxonomic group, with the magnitude of difference between empirical and expected PD increasing with elevation for all passerines and suboscines (Fig. 2). In contrast, empirical elevational variation in FD was no different than expected based on random selection for each taxonomic group (Fig. 2). When decomposed into constituent functional components, only diet diversity and size diversity for oscines deviated from expectations based on random selection, declining faster than expected in each case (Fig. 3).

Differences in elevation-specific species richness between suborders were not significant (Paired-t = −1.97, DF = 12, P = 0.072). In contrast, suboscines had greater elevation-specific PD (Paired-t = −13.66, DF = 12, P < 0.001), whereas oscines had greater elevation-specific FD (Paired-t = 7.07, DF = 12, P < 0.001). In addition, oscines exhibited greater diet (Paired-t = 19.24, DF = 12, P < 0.001) and size (Paired-t = 5.50, DF = 12, P < 0.001) diversity, whereas suboscines exhibited greater foraging location diversity (Paired-t = −8.91, DF = 12, P < 0.001). The suborders did not differ in foraging strategy diversity in an elevation-specific manner (Paired-t = 1.04, DF = 12, P = 0.316).

For species richness and FD, linear (b_1_) and non-linear (b_2_) parameters from ordinary polynomial regression, and the coefficients that represent independent contributions of the linear (b^*^_1_) and non-linear (b^*^_2_) components from orthogonal polynomial regression, were all significantly different between oscines and suboscines (Fig. 4). In contrast, only the independent contribution of the linear component (b^*^_1_) from orthogonal polynomial regression was significantly different between suborders for PD. Elevational relationships for foraging strategy diversity and foraging location diversity were not significantly different between suborders (Supplementary material Fig. S1). Conversely, the slopes for size diversity and the linear and quadratic components for diet diversity were different between suborders, indicating that these two components of functional diversity are most responsible for differences between suboscines and oscines in elevational gradients of FD.

**Figure 4.**
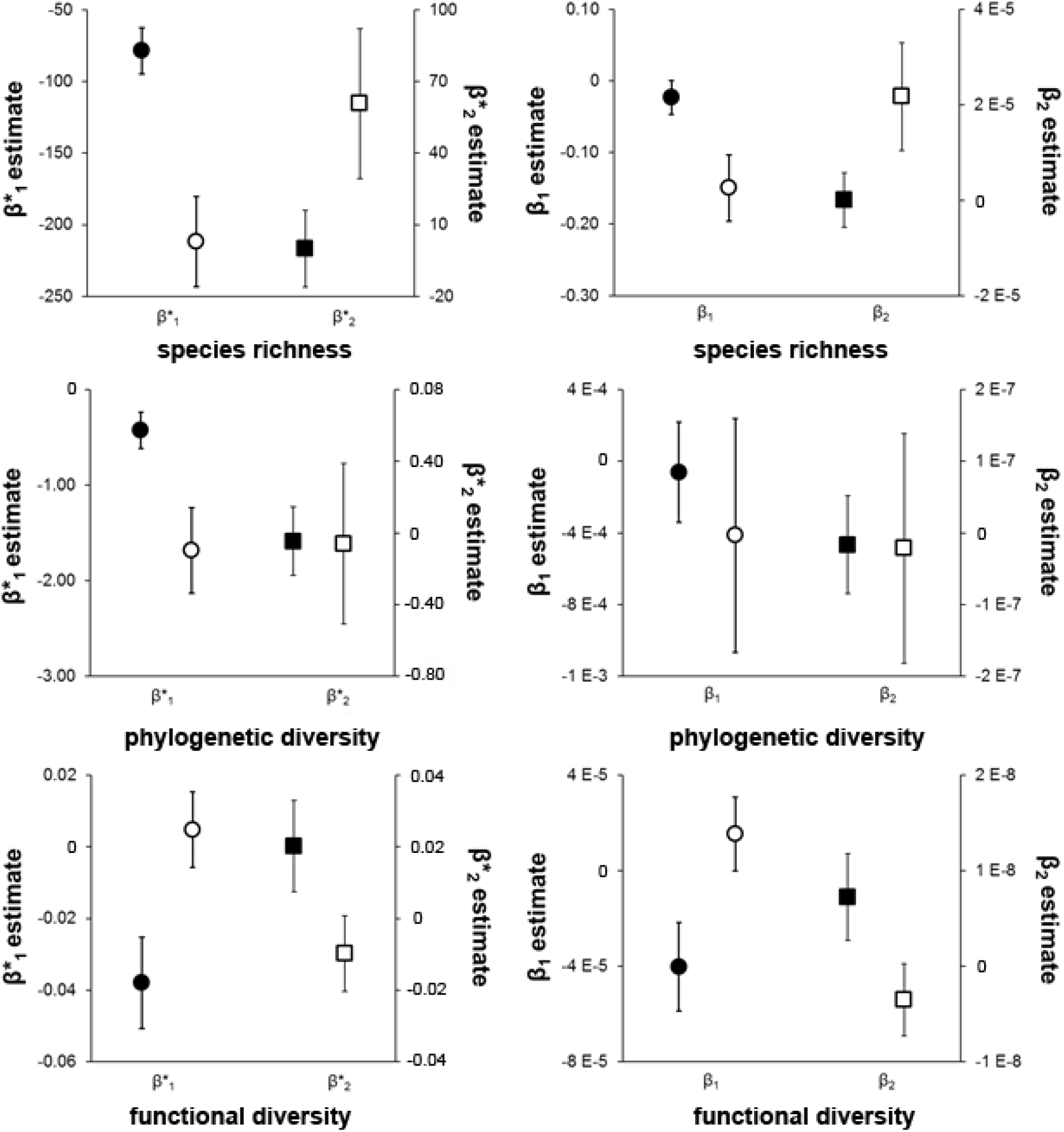
Comparisons of parameter estimates for elevational gradients of species richness, phylogenetic diversity (PD), and functional diversity (FD) for oscines (solid symbols) and suboscines (open symbols). Error bars are ± 2 SE. Circles represent the estimated values for the linear component (b*1) from orthogonal polynomial regression and the slope (b1) from ordinary polynomial regression. Squares represent the estimated values for the quadratic component (b*2) from orthogonal polynomial regression and the quadratic term (b2) from ordinary polynomial regression. Parameter estimates for which confidence intervals do not overlap were considered to be significantly different.

## DISCUSSION

Biodiversity declined with elevation for the taxonomic and phylogenetic dimensions, but elevational gradients of FD were contingent on taxon and the functional traits (i.e., diet, foraging location, foraging strategy, body size) used to characterize it (Fig. 1–3). Any dimension or functional component of biodiversity that exhibited a negative relationship with elevation was positively correlated with species richness (Table 2). Nonetheless, these relationships exhibited multiple forms: linear, saturating at high elevations (e.g. FD for oscines), or saturating at low elevations (e.g. diet for suboscines). All deviations from expectations based on random sampling were associated with diversity declining with elevation faster than expected given empirical species richness (PD for all groups, diet and size diversity for oscines). In addition to the form of elevational gradients differing between oscines and suboscines (Fig. 1–3), PD of suboscines was consistently greater than that of oscines throughout the gradient, whereas FD of oscines was consistently greater than that of suboscines throughout the gradient. In general, these results are related to differences between the suborders in historical biogeography and level of conservatism of ancestral niches, patterns that would have been obscured had we only considered passerines as a whole.

### Species richness

Differences between suboscines and oscines in the elevational gradients of species richness at Manu reflect their distinct evolutionary origins. Suboscines are more species-rich than are oscines in the lowlands, but oscines are more species-rich than are suboscines above 1750 m. This pattern is more pronounced when comparisons are based on the proportion of passerine species from each lineage (Fig. 1d). Before the Miocene, South America was largely lowland and tropical, resulting in the development of warm-weather adaptations early in the evolutionary history of suboscines (Fedducia 1999). Oscines generally have higher activity levels, higher metabolic rates, and greater cold tolerance than do suboscines (Feduccia 1999, Swanson and Bozinovic 2011), likely contributing to oscines being more species-rich than suboscines in the colder climates of high elevations.

Additionally, many oscine groups diversified in temperate areas (Barker et al. 2004), likely resulting in adaptations to deal with temporal or spatial variation in biotic (e.g. resource abundance and diversity) or abiotic conditions (Ricklefs 2002). Finally, the Andean uplift coincided with tanager diversification into one of the most ecologically diverse avian families (Fjeldsa and Rahbek 2006, Burns et al. 2014), which suggests why they largely drive patterns of oscine species richness at Manu (Fig. 5). Importantly, vertebrates of tropical origin exhibit greater thermal niche conservatism than do vertebrates of temperate origin (Cadena et al. 2012), suggesting that climatic niche conservatism may have been a greater constraint for tropical suboscines adapting to colder, high-elevation environments as the Andes rose than it was for the temperate oscines adapting to warmer, low-elevation environments. In summary, biogeographic history, evolutionary contingencies, and climatic niche conservatism within each suborder contributed to the elevational gradients of species richness in oscines and suboscines.

**Figure 5.**
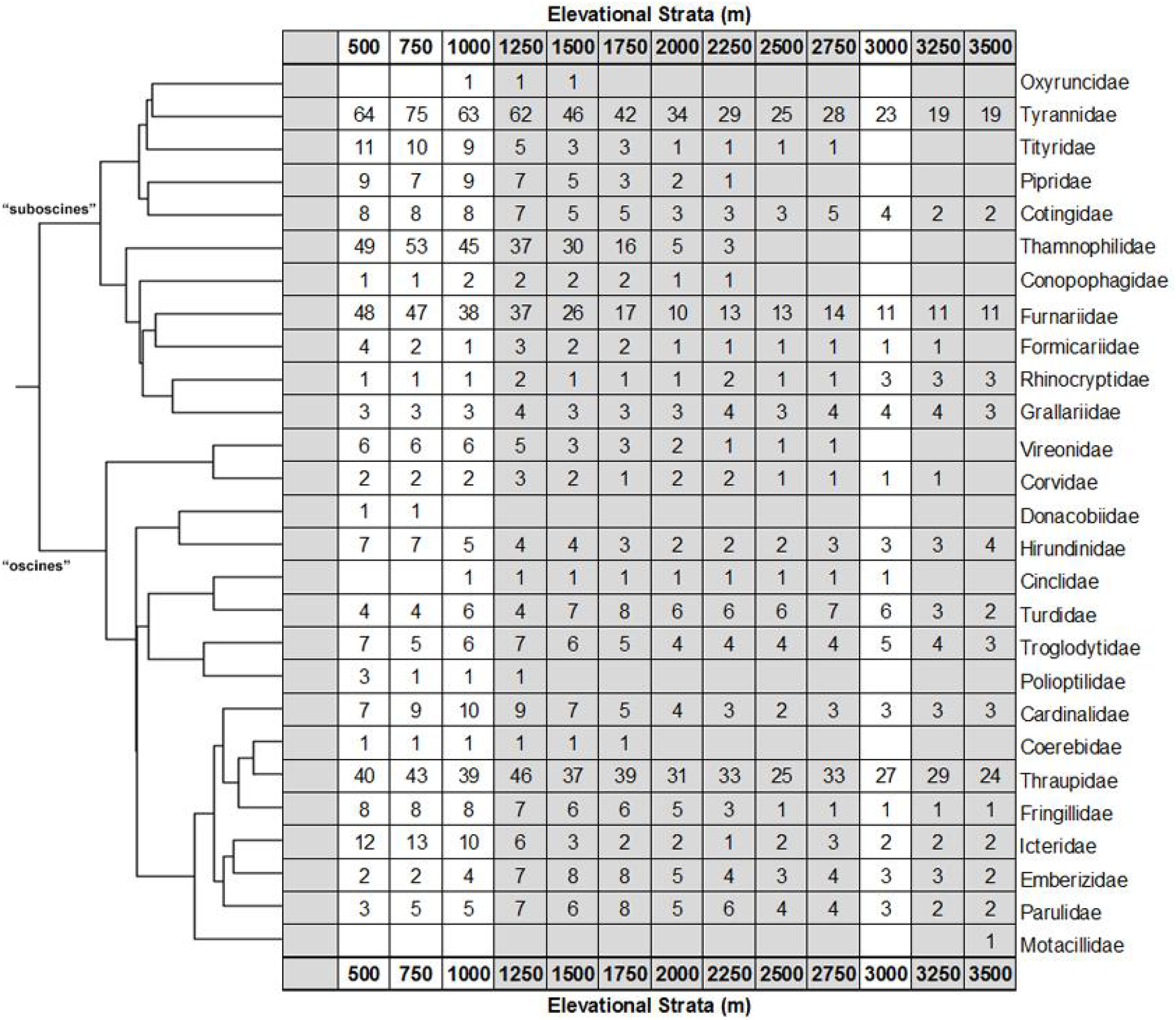
Elevational variation in species richness for each passerine family at Manu based on the phylogeny of Jetz et al. (2012). Alternating shaded regions of the graphic correspond to elevationally defined forest types: lowland rainforest, montane rainforest, cloud forest, elfin forest, and mixed elfin forest-grassland.

### Phylogenetic biodiversity

PD decreased along the elevational gradient for each taxonomic group (Fig. 2), consistent with the results based on global patterns of avian communities along mountain gradients (Montano-Centellas et al. 2019) and for only frugivorous birds at Manu (Dehling et al. 2014). This gradient of PD is robust with respect to metric choice as the study on frugivores used Faith’s PD, which sums all branch lengths for a phylogeny of the community, whereas we used a dispersion metric (Rao’s Q) transformed into a numbers equivalent. The elevational decline of passerine PD, as measured by Rao’s Q, can be attributed to the differential loss of species from particular families and the differential loss of entire families with increasing elevation. These changes in familial richness are consistent with greater thermal niche conservatism and associated specialization of suboscines on lowland habitats, whereas oscine lineages typically occur along most of the gradient, with relatively small changes in species richness between vegetation types (Fig. 5).

The total number of passerine families declined from 24 in lowland rainforest to 15 in the mixed elfin forest and montane grasslands (Fig. 5), and the number of relatively species-rich families (10 or more species) declined from 6 (2 oscine and 4 suboscine) to 3 (1 oscine and 2 suboscine). The suboscines lost a greater proportion of families (45%) with increasing elevation than did oscines (28%). Tanagers (Thraupidae, 29 species), tyrant flycatchers (Tyrannidae, 19 species), and ovenbirds (Furnariidae, 11 species) are the only families represented by > 4 species above 3000 m. These three families comprise 65% of all passerine species at these high elevations. In concert, these changes resulted in the tanagers accounting for > 54% of oscine species and > 29% of all passerine species at elevations above 3000 m. Despite representing multiple radiations before their arrival in South America and multiple distinct colonization events, the dominance of the oscines by tanagers resulted in consistently lower PD for oscines than for suboscines along the gradient (Fig. 2). This dominance may have occurred because tanagers diversified from the Andean highlands down to adjacent lowlands, and did so during the Andean uplift (Fjeldsa and Rahbek 2006). As elevation increases, this increased “clustering” of species that represent a few dominant lineages (i.e. tanagers, flycatchers, and ovenbirds) accounts for the negative relationship of PD with elevation for each taxonomic group (Fig. 2).

The increasing dominance of a few families (e.g., Furnariidae, Thraupidae, Tyrannidae) with increasing elevation resulted in steeper elevational gradients of PD than expected based on random selection for each species group (Fig. 2). However, the relationships of the empirical and expected gradients differed between suborders. For suboscines, empirical and expected PD were similar at low elevations, indicating that empirical community composition did not differ from that associated with a random sample of oscine lineages. However, the differential loss of species from different families, particularly the reduction in richness from 49 at 500 m to 0 at 2500 m of the antbirds (Thamnophilidae), resulted in a greater decrease in PD than expected by chance. In contrast, empirical PD for oscines was greater than mean expectations at low elevations, indicating a relatively even representation of oscine lineages in lowland forests, and was less than mean expectations at high elevations due to the dominance of tanagers in those assemblages (Fig. 2). These gradients of PD arise because the retention of historical ecological functions limits the elevational distributions of some lineages (e.g. conservatism of thermal niches in clades of tropical origin; Cadena et al. 2012).

### Functional biodiversity

Variation in FD based on all functional attributes was not associated with elevation for all passerines or suboscines (Fig. 2), contrasting with the negative elevational relationship of FD for oscines (Fig. 2) and for frugivorous birds at Manu based on morphometric traits associated with feeding, flight, and bipedal locomotion (Dehling et al. 2014). A higher proportion of oscines (59%) at Manu are frugivorous than are suboscines (32%), which likely explains both similarities in the gradients for all frugivorous birds and oscines, as well as the origin of the differences between all frugivorous birds and suboscines. Importantly, diet diversity decreased for both suboscines and oscines (Fig. 3), indicating that the reduction in the abundance and diversity of dietary resources may mold patterns of FD within clades that represent multiple foraging guilds (i.e. oscines and suboscines) as well as within foraging guilds that comprise many lineages (i.e. the avian frugivores of Dehling et al. 2014).

The use of only morphometric traits may have contributed to the decrease in FD for frugivorous birds with increasing elevation (Dehling et al. 2104), as the only functional component based on morphometrics in our study, body size, evinced a negative relationship with elevation for all passerines and for oscines (Fig. 3). Importantly, these results do not indicate that bird morphometric traits are smaller at high elevations, only that these traits are more similar (i.e., occupy a smaller volume of morphometric space) with increasing elevation. Results from both studies suggest that environmental constraints associated with harsh environments at higher elevation may decrease diversity in bird morphology and size. Although body size diversity of suboscines did not exhibit a consistent elevational pattern, the body size diversity of oscines and suboscines converged with increasing elevation (Fig. 3). Oscines are larger than suboscines at low elevations, but the lineages are similar in size from ~1500 m to the top of the Andes (Fig. S2). Convergence in body size and body size diversity between lineages suggests the existence of strong constraints at higher elevations associated with thermoregulation and resource availability.

Despite the absence of an elevational gradient of FD for all passerines, each functional component of FD did exhibit a significant response to elevation. Diet and size diversities decreased with elevation, whereas foraging location and foraging strategy diversities increased with elevation (Fig. 3). These divergent responses highlight the importance of separate consideration of functional components that represent distinct niche axes. At Manu, divergent responses of functional components also occurred for bats (Cisneros et al. 2014) but did not occur for rodents (Dreiss et al. 2015). For passerines and bats, foraging strategy and foraging location diversity increased with elevation, whereas diet and size diversity decreased with elevation, indicating that similar ecological mechanisms (e.g. physiological constraints associated with temperature, reduction in functional redundancy associated with decreasing resource abundance and diversity) may shape responses of volant vertebrates to environmental variation associated with elevation.

Diet diversity decreased with elevation for each lineage of passerines (Fig. 3); however, diet diversity in oscines varied little in the cloud forest and above, and diet diversity of suboscines peaked in rain forest. Regardless of elevation, oscines had greater diet diversity than did suboscines (Fig. 3). These differences in diet diversity are largely related to physiological and morphological differences between lineages. For example, oscines are more active predators, whereas suboscines are more likely to employ a sit-and-search strategy to acquire prey (Ricklefs 2002, Swanson and Bozinovic 2011). Differences in prey capture and activity levels allow oscines to dominate in the canopy and open spaces, whereas suboscines perform better in the subcanopy (Ricklefs 2002). These ecological differences are reflected in differences in morphology, as suboscines are generally less mobile and less agile than are oscines, with shorter wings, legs, toes, and tails (Ricklefs and Travis 1980, Rickefs 2002). These traits also make it more difficult for suboscines to pursue prey through the canopy (Ricklefs 2002). Temperature, net primary productivity, and insect abundance decrease with elevation in the Andes (Garibaldi et al. 2011). As a result, the richness of avian insectivores is disproportionately lost with increasing elevation (Terborgh 1977, Jankowski et al. 2013), which should affect suboscines more than oscines, and contribute to oscine dominance at high elevations.

Foraging location and strategy diversities increased with elevation for all passerines, with suboscine foraging location diversity and oscine foraging strategy diversity increasing with elevation (Fig. 3). Suboscines dominate the subcanopy of rainforests; however, the subcanopy becomes indistinguishable from the understory as elevation increases. This may result in differential loss of suboscines that use sit-and-search strategies in the subcanopy from high-elevation assemblages (Jankowski et al. 2013). Loss of species with the same foraging location and foraging strategy attributes results in greater diversity at higher elevations for each of those functional components. Importantly, flycatchers (Tyrannidae) are more like oscines in form and habit than they are like other suboscines and do well both in open areas and in the canopy (Ricklefs 2002). The ecological differences between the Tyrannidae and Furnariidae (both suboscines) result in greater foraging location diversity for suboscines than for oscines throughout the gradient (Fig. 3). In general, suboscines have longer, thinner beaks than do oscines, making them better suited to sit-and-search foraging tactics in the subcanopy and to capturing larger insects (Ricklefs and Travis 1980, Ricklefs 2002). Nonetheless, these adaptations limit the ability of suboscines to process fruit, with some notable exceptions (e.g. the Cotingidae). In addition, oscines are typically better able to respond to temporal variation in resource availability because they are more euryphagic (Ricklefs 2002), an important adaptation to maintain viable populations in low productivity environments.

## Conclusions

Explicit consideration of history – including historical biogeography, historical contingency, and conservatism of functional traits – is necessary for a more nuanced understanding of the ecological patterns of species richness as well as of phylogenetic and functional biodiversity. Most studies that evaluate spatial patterns of PD and FD involve trait conservatism and simple evolutionary histories that result in congruence between these dimensions (e.g. Cadotte et al. 2009, Devictor et al. 2010, Flynn et al. 2011, Meynard et al. 2011, Cisneros et al. 2014, Dehling et al. 2014, Dreiss et al. 2015). In contrast, the passerines at Manu provide an example of the decoupling of functional biodiversity from phylogenetic biodiversity. For Manu passerines, elevational gradients of biodiversity comprise multiple clades, each representing independent radiations driven by different environmental factors (e.g. suboscines in Amazonian lowlands, oscines in colder climes of North America, and tanagers evolving in situ in the Andes). Passerine communities along an elevational gradient in the Andes likely are molded by a combination of ecological factors, most importantly, physiological constraints associated with temperature, and a reduction in functional redundancy associated with decreasing forest stratification and decreasing resource abundance and diversity.

Effective generalizations for spatial patterns of multiple dimensions of biodiversity have been stymied by the relative dearth of studies that quantify gradients for multiple dimensions, resulting in poor coverage from taxonomic and geographic perspectives. In general, biodiversity gradients for any species-rich biota likely are affected by the niche characteristics of constituent species, the nature of the environmental gradient, the strength of niche conservatism in the biota, and biogeographic or historical contingencies associated with the evolutionary history of the fauna and region. The relative importance of these mechanisms will determine the form of spatial gradients and relationships among dimensions. For example, bats (Cisneros et al. 2014), rodents (Dreiss et al. 2015), and passerines (this manuscript) from Manu have similar elevational gradients of species richness due to the pervasive effects of productivity on species richness, but gradients of PD and FD are taxon-specific due to differences between groups in how evolutionary histories, niche conservatism, and ecological functions performed by each taxon combine to shape elevational patterns.

## ACKNOWLEDGMENTS

We are especially grateful to B. Patterson for his insights and assistance, as well as to J. Bates for helpful comments. Funding for the synthetic portion of this project was provided by a National Science Foundation (NSF) Grant to S. Andelman and J. Parrish entitled “The Dimensions of Biodiversity Distributed Graduate Seminar” (DEB-1050680). KRB was supported by an NSF GRFP grant (DGE-0753455) and an NSF NRT-IGE grant (#1545458) awarded to M. Rubega. LMC was supported by a Multicultural Fellowship from the Graduate School at the University of Connecticut. MRW and SJP were supported by the Centre for Environmental Sciences and Engineering at the University of Connecticut and by NSF grants (DEB-0620910, DEB-1239764, DEB-1546686 and DEB-1831952) to the Institute of Tropical Ecosystem Studies, University of Puerto Rico, and the International Institute of Tropical Forestry as part of the Long-Term Ecological Research Program in the Luquillo Experimental Forest. The authors declare no conflicts of interest.

## BIOSKETCH

**Kevin R. Burgio** is broadly interested in the processes that form, and limit, where species are distributed, and their roles are in communities and ecosystems. He uses an integrative approach to examine historical ecology, community assembly, the effects of climate change on communities, and extinction (webpage: https://kevinburgio.com/).

Author contributions: All authors conceived of the ideas; KRB compiled the trait data; KRB and SJP conducted all analyses. KRB led and all authors contributed to the writing of the manuscript.

