## Supplemental Information for "Dimensions of passerine biodiversity along an elevational gradient: a nexus for historical biogeography and contemporary ecology"

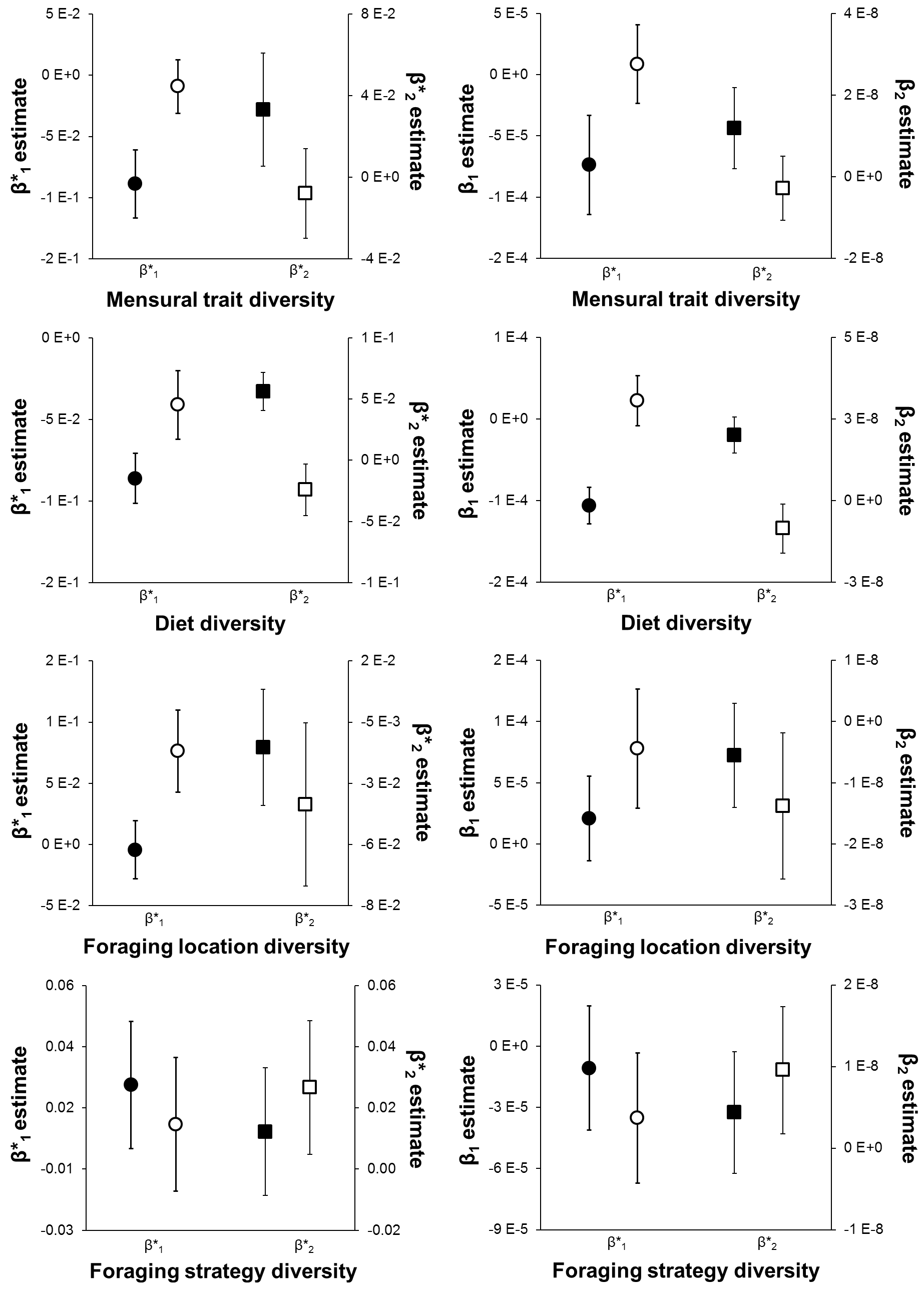

Figure S1. Comparisons of parameter estimates for elevational gradients of functional diversity based on consideration of each functional component separately for oscines (solid symbols) and suboscines (open symbols). Error bars are ± 2 SE. Circles represent the estimates for the linear component (b*_1_) from orthogonal polynomial regression and the slope (b_1_) from ordinary polynomial regression. Squares represent the estimates for the quadratic component (b*_2_) from orthogonal polynomial regression and the quadratic term (b_2_) from ordinary polynomial regression. Parameter estimates for which confidence intervals do not overlap were considered to be significantly different.

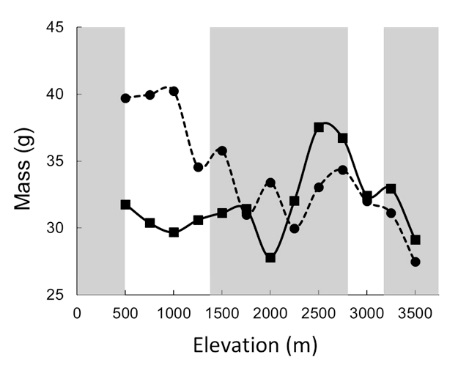

Figure S2. Mean mass at each elevational stratum for oscines (circles and dashed line) and for suboscines (squares and solid line).

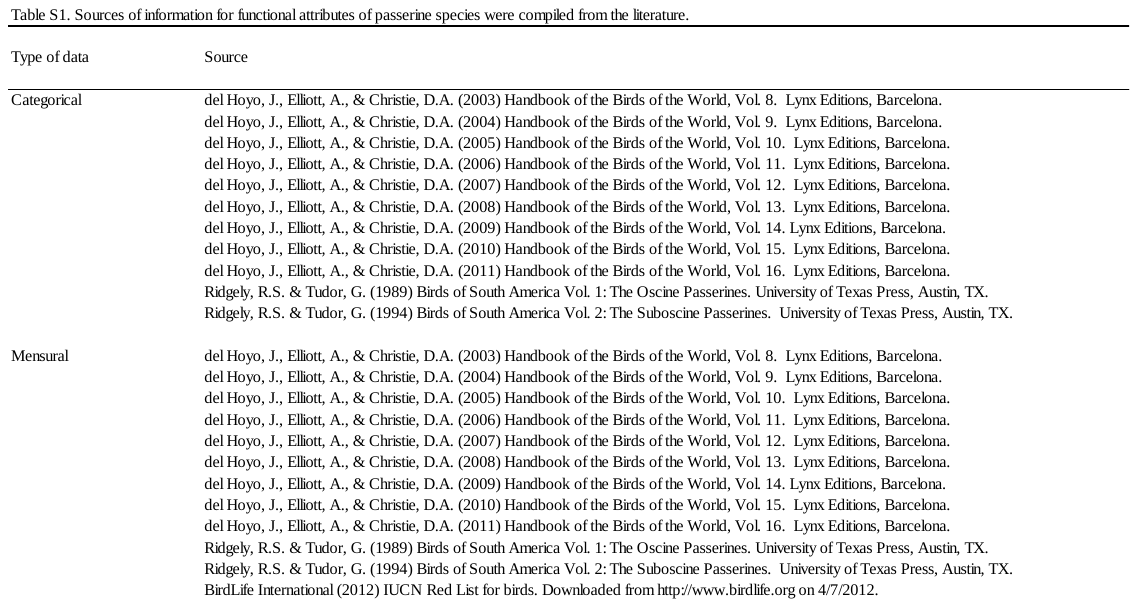

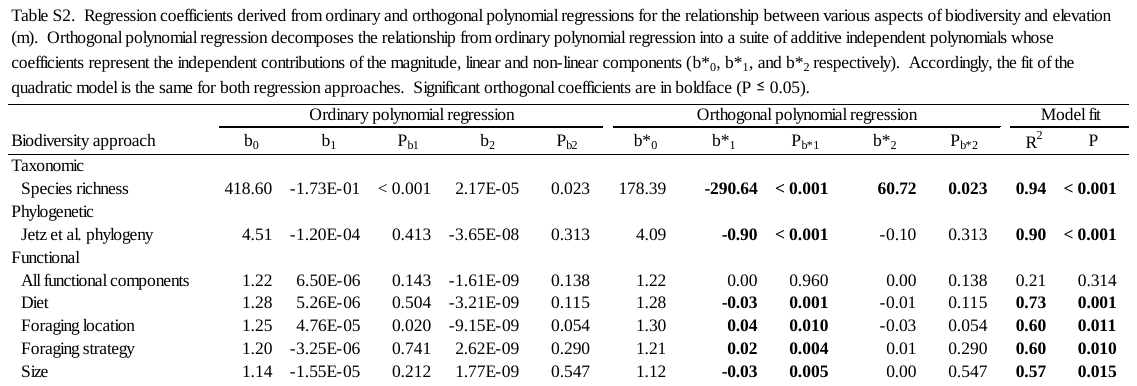

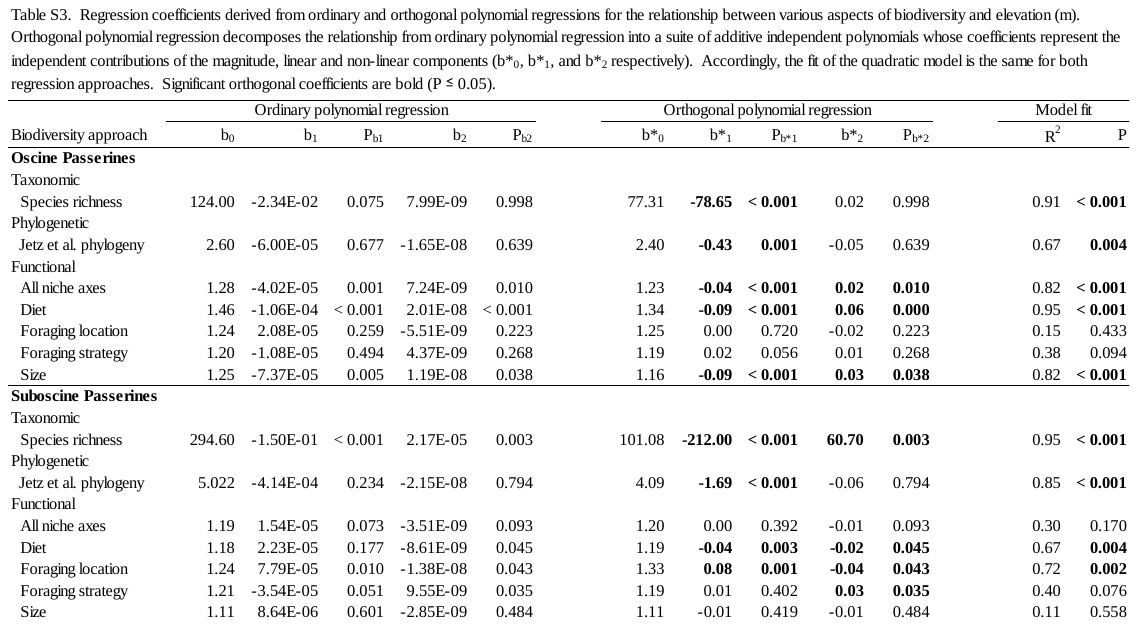
